# Perception affects the brain’s metabolic response to sensory stimulation

**DOI:** 10.1101/2021.09.17.460759

**Authors:** Mauro DiNuzzo, Silvia Mangia, Marta Moraschi, Daniele Mascali, Gisela E. Hagberg, Federico Giove

**Affiliations:** Museo Storico della Fisica e Centro Studi e Ricerche Enrico Fermi, 00184 Rome, Italy; Center for Magnetic Resonance Research, Department of Radiology, University of Minnesota, Minneapolis, MN 55455, USA; Unità Operativa di Radioterapia Oncologica, Università Campus Bio-Medico, 00128 Rome, Italy; Dipartimento di Neuroscienze, Imaging e Scienze Cliniche, Università Gabriele D’Annunzio, 66100 Chieti, Italy; High–Field Magnetic Resonance, Max Planck Institute for Biological Cybernetics and Biomedical Magnetic Resonance, University Hospital Tübingen, 72076 Tübingen, Germany; Fondazione Santa Lucia IRCCS, 00179 Rome, Italy

**Keywords:** central visual system, visual perception, isoluminant chromatic flickering, human brain metabolism, lactate, BOLD fMRI, single voxel ^1^H-fMRS

## Abstract

Processing of incoming sensory stimulation triggers an increase of cerebral perfusion and blood oxygenation (neurovascular response) as well as an alteration of the metabolic neurochemical profile (neurometabolic response). Here we show that perceived and unperceived isoluminant chromatic flickering stimuli designed to have similar neurovascular responses as measured by blood oxygenation level dependent functional MRI (BOLD-fMRI) in primary visual cortex (V1) have markedly different neurometabolic responses as measured by functional MRS. In particular, a significant regional buildup of lactate, an index of aerobic glycolysis, and glutamate, an index of malate-aspartate shuttle, occurred in V1 only when the flickering is perceived, without any relation with behavioral or physiological variables. Wheras the BOLD-fMRI signal in V1, a proxy for input to V1, was insensitive to flickering perception by design, the BOLD-fMRI signal in secondary visual areas was larger during perceived than unperceived flickering indicating increased output from V1. These results indicate that the upregulation of energy metabolism induced by visual stimulation depends on the type of information processing taking place in V1, and that 1H-fMRS provides unique information about local input/output balance that is not measured by BOLD-fMRI.

**Significance statement:** Visual perception has a measurable metabolic effect in the primary visual cortex (V1). Here we show that the upregulation of energy metabolism induced by isoluminant chromatic flickering depends on subjective visual perception. Within V1, perceived and unperceived stimuli that are contrast-matched to elicit similar blood-oxygenation level-dependent fMRI response are associated with clearly distinct neurochemical profiles. Specifically, regional accumulations of lactate (an index of aerobic glycolysis) and glutamate (an index of malate-aspartate shuttle) only develop during perceived stimuli, together with a larger activation of secondary visual areas. Our results imply a dissociation between metabolic and functional response, and indicate that that the upregulation of energy metabolism induced by visual stimulation depends on the type of information processing taking place in V1.

## Introduction

In the brain, sensory stimulation is associated with a substantial increase of regional functional hyperaemia (i.e. cerebral blood flow, CBF) as well as energy metabolism of glucose, the main cerebral energy substrate (1). The metabolic response to stimulation includes an oxidative component, as measured by the cerebral metabolic rate of oxygen (CMRO_2_), and a non-oxidative component, as reflected by lactate accumulation (2). Cortical lactate levels have been shown to increase during visual stimulation, simultaneously to the acceleration of the malate-aspartate shuttle, a process termed aerobic glycolysis (i.e. lactate production independent of oxygen availability) (3–11). Despite intense research, the neurophysiological mechanisms underlying the up-regulation of glycolytic metabolism of glucose are still largely unknown (12). Most importantly, the impact of information processing to the metabolic response of the cerebral cortex to sensory stimulation has not been fully investigated thus far. In particular, nothing is known about the modulatory effect exerted by the perception of different stimuli on regional brain energy metabolism.

Sensory perception is thought to rely on the complex interplay of neural circuits that process information in a cortical layer-and area-mediated manner involving thalamo-cortical, intracortical, cortico-cortical and cortico-thalamic feedforward/feedback loops (13). Sensory stimuli transduced by sensory organs reach specific thalamic nuclei that relay information to primary sensory cortices, which in turn filter and eventually transmit information to secondary sensory areas (14). These transactions are dependent on the particular features of different incoming stimuli, thus it is possible that the relevant neurovascular and neurometabolic counterparts are correspondingly distinct (15).

The thalamic lateral geniculate nucleus (LGN) mediates visual stimuli with temporal frequencies at least up to 90 Hz to the layer IV of V1 (16–21), which in turn relays to output layers II/III and V where temporal filtering occurs (22), consistent with the notion that visual perception requires the activation of visual areas downstream V1 (i.e. secondary visual cortices). In agreement with these arguments, it has been repeatedly reported that invisible visual flickering is still able to activate V1 even without any perceptual effects (23), as revealed by in vivo electrophysiology in non-human primates (16) as well as behavioral evidence (24) and BOLD fMRI (25) in humans. High (30 Hz) frequency visual stimulation has been found to selectively suppress multi-unit activity (MUA) in cat V1 as compared to low frequency (4 Hz) visual stimulation (26). Importantly, local field potentials (LFPs) and tissue oxygen response, which directly contribute to the generation of the BOLD signals (27), were preserved at both frequencies.

In the present study, we combined blood-oxygenation level dependent (BOLD) functional magnetic resonance imaging (fMRI) and proton functional magnetic resonance spectroscopy (1H-fMRS) in humans and exploited the well known effect of temporal frequency on visual perception. Specifically, we examined the functional and metabolic responses of the primary visual cortex (V1) to perceived or unperceived isoluminant chromatic flickering stimulations obtained by using temporal frequency either below (7.5 Hz; PF, perceived flickering) or above (30 Hz; UF, unperceived flickering) the critical flicker fusion (CFF) threshold of ∼15 Hz for rod-mediated vision (28). Based on experimental evidence and metabolic modeling, we have previously proposed that the local input-output balance between neuronal synaptic/spiking (or subthreshold/suprathreshold) activity is a primary determinant in the up-regulation of aerobic glycolysis (29–31). We thus hypothesized that the loss of visual perception is accompanied by fundamental changes in the metabolic responses of human V1.

## Results

### Subjects perception of the visual stimuli

To achieve perceptual isoluminance between green and red color (necessary for loss of perception at 30 Hz), we adjusted the brightness of the green color for each individual subject, which was remarkably similar across subjects (green/red brightness ratio 71.9±1.2%, range 70.1% to 73.5%; see Table 1). After this procedure, 100% of the subjects confirmed that their perception of the 30 Hz frequency stimulus steadiness was equivalent to the resting condition. Overall, the subject’s perception was a gray/colored checkerboard that in the colored squares showed either a fast green and red alternation during PF epochs, or a static yellow during UF epochs (Movie S1). As a further confirmation, while in the scanner the subjects were unable to distinguish the 30 Hz red-green flickering checkerboard (used in the actual experiments) from a color-matched static yellow checkerboard (used for testing only). Specifically, the perception of the steady yellow color versus the 30 Hz red-green flickering was indistinguishable, as assessed by asking the subjects to guess the origin of the stimulus for 10 consecutive trials (average of correct responses 52±16%, not different from chance level, p=0.62). All subjects reported to distinctly perceive the green and red color when the checkerboard was flickering at 7.5 Hz. None of the subject perceived the intrinsic flickering of the screen due to the refresh rate (60 Hz).

**Table 1.** Demographics and study parameters.

| Count | Gender | Age | fMRI |  | 1H-fMRS |  |  |
| --- | --- | --- | --- | --- | --- | --- | --- |
|  |  |  | Calibration | Run | Voxel | Runs | Water Linewidth (Hz) |
| 1 | M | 35 | 72.5 | 1 | L | 1/2 | 6.8 |
| 2 | F | 32 | 72.5 | 2 | L | 2/1 | 8.3 |
| 3 | M | 29 | 73.5 | 1 | R | 1/- | 7.5 |
| 4 | F | 24 | 71.8 | 2 | R | 2/1 | 7.7 |
| 5 | M | 26 | 72.1 | 1 | L | 1/2 | 6.7 |
| 6 | F | 27 | 73.5 | 2 | L | 2/1 | 6.7 |
| 7 | M | 28 | 70.1 | 1 | L | 1/2 | 5.7 |
| 8 | M | 20 | 72.1 | 1 | R | 2/1 | 7.5 |
| 9 | F | 26 | 70.1 | 2 | L | 1/2 | 6.6 |
| 10 | F | 28 | 72.1 | 1 | R | 2/1 | 6.8 |
| 11 | F | 20 | 70.1 | 2 | R | 1/2 | 7.5 |
| 12 | F | 20 | 73.5 | 1 | L | 1/2 | 7.9 |
| 13 | F | 20 | 71.8 | 1 | R | 2/1 | 7.8 |
| 14 | M | 21 | 73.5 | 2 | L | 2/1 | 7.4 |
| 15 | M | 21 | 71.8 | 1 | L | 2/1 | 7.6 |
| 16 | F | 26 | 70.1 | 2 | R | 2/1 | 7.5 |
| 17 | M | 20 | 70.1 | 1 | R | 2/1 | 7.0 |
| 18 | F | 26 | 72.8 | 2 | R | 1/2 | 6.3 |
| 19 | M | 25 | 71.8 | 1 | L | 2/1 | 7.1 |

Calibration refers to the % brightness adjustment for the green color relative to the brightness of the red color. A dash indicates that the relevant measurement has not been performed due to technical reasons.

### Subjects attention

To examine the possible modulation of the BOLD signal by attention (32), we measured pupillometry and task-performance data during all experiments. All subjects maintained gaze on the fixation cross during the whole epochs (Figure 1A), with no difference in average gaze location between the different stimulations (Figure 1B,C and Figure 1 – figure supplement 1^1^). Pupil diameter, an index of the noradrenergic tone (33), was fairly stable across conditions (Figure 1D,E; Figure 1 – figure supplement 2 and Movie S1), indicating that the modulation of perception by noradrenaline (34) was minimal in our experimental conditions.

**Figure 1.**
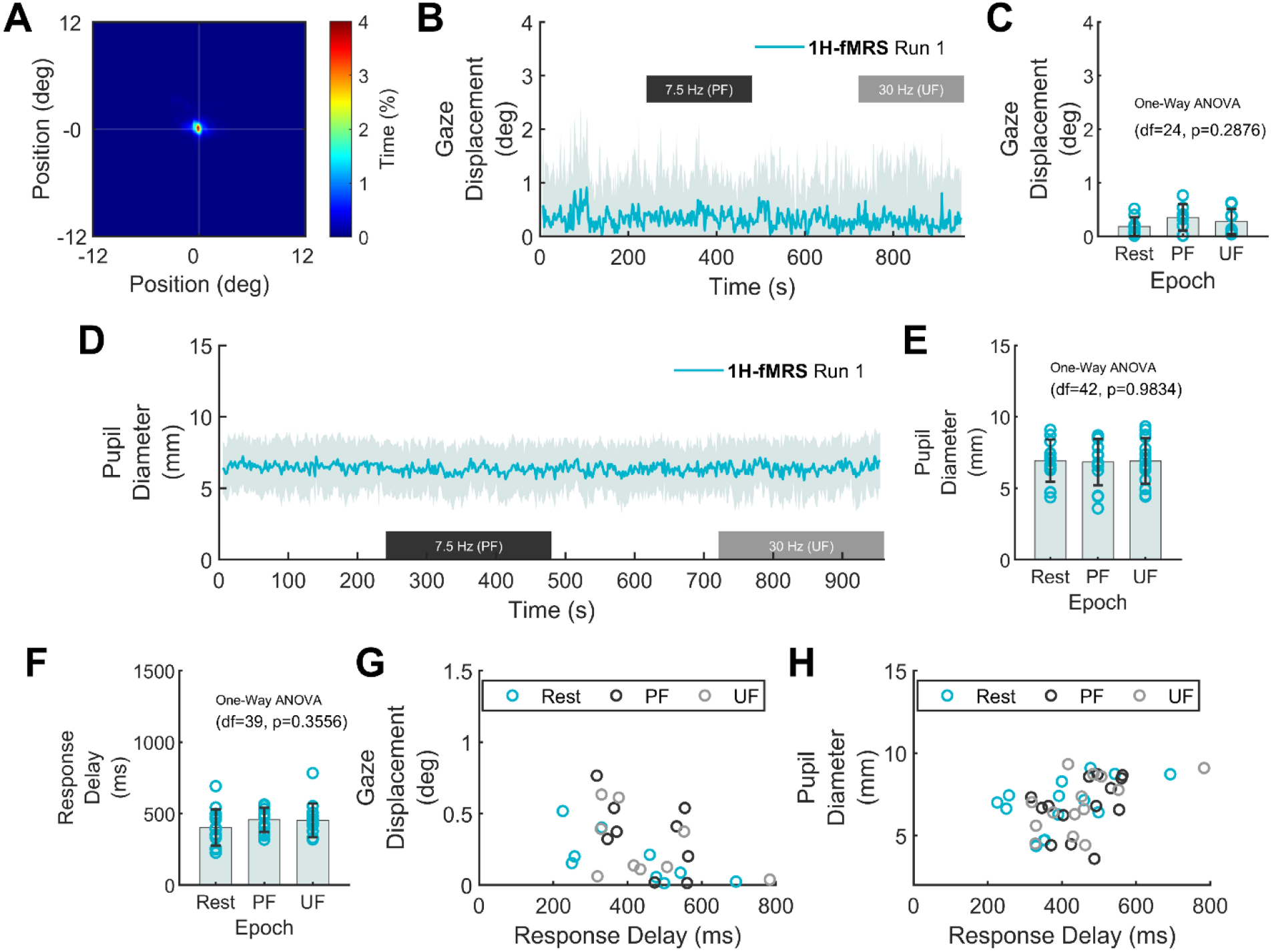
Subjects perception and attention during visual stimulation. (A) Average heatmap of eyes position (across subjects) during a representative session (1H-fMRS Run 1). (B) Stability of mean gaze displacement from the fixation point. (C) Gaze displacement was not different across conditions (One-Way ANOVA, p=0.29). Error bars correspond to SD. (D) Average pupil diameter (across-subjects) during a representative session (1H-fMRS Run 1). (E) Mean pupil diameter was not statistically different across conditions (One-Way ANOVA, p=0.98). Error bars correspond to SD. (F) Task performance in terms of response delay was not statistically different across conditions (One-Way ANOVA, p=0.36). Error bars correspond to SD. (G) There was no correlation between task performance and mean gaze displacement (r^2^<0.06, p>0.47). (H) There was no correlation between task performance and mean pupil diameter (r^2^<0.08, p>0.49).

The hit/miss ratio for the cross rotation task during the stimulation protocol was essentially 1.0, as expected due to the simplicity of the task, for the entire duration of the experiment. In particular, the delay of the response was not statistically different for rest, PF and UF epochs (ranging, on average, between 360 ms and 460 ms), both during fMRI (One-Way ANOVA, p=0.09 and p=0.77 for run 1 and 2, respectively) and 1H-fMRS (One-Way ANOVA, p=0.39 and p=0.51 for run 1 and 2, respectively), confirming high and similar levels of subject’s attention across conditions (Figure 1F and Figure 1 – figure supplement 3). There was no significant correlation between task performance and mean pupil diameter or gaze displacement (p>0.21 and p>0.34, respectively; Figure 1G,H and Figure 1 – figure supplement 3). Finally, in-scanner head motion during fMRI scans was minimal and no significantly different for all subjects across epochs (mean framewise displacement 0.25±0.12 mm for rest, 0.24±0.14 mm for PF, 0.23±0.08 mm for UF; One-Way ANOVA, p=0.77). Overall, behavioral and physiological variables associated with attentional load were maintained at considerably constant levels in all subjects.

### Similar BOLD responses in V1 to perceived and unperceived flickering

To achieve the same BOLD response in V1 during PF and UF, we reduced the stimulation contrast for the 7.5 Hz condition to 75% relative to the 30 Hz condition (Figure 2A). As expected, we found that the average BOLD timecourse (Figure 2B) as well as the change in the subject-matched spectroscopic VOI (on average consisting of 47±9% of BA17, 21±12% of BA18, and 16±9% of BA19; see Figure 2 – figure supplement 1), was similar between the two conditions (0.44±0.30% for PF versus 0.41±0.25% for UF, paired two-sample t-test, p=0.71) (Figure 2C). The fMRI activations to PF and UF both peaked in V1 and distinctly spanned bilaterally in secondary visual areas (Figure 2D,E, one-sample t-test, FDR-corrected at cluster level, q<0.05, voxel level p<0.001).

**Figure 2.**
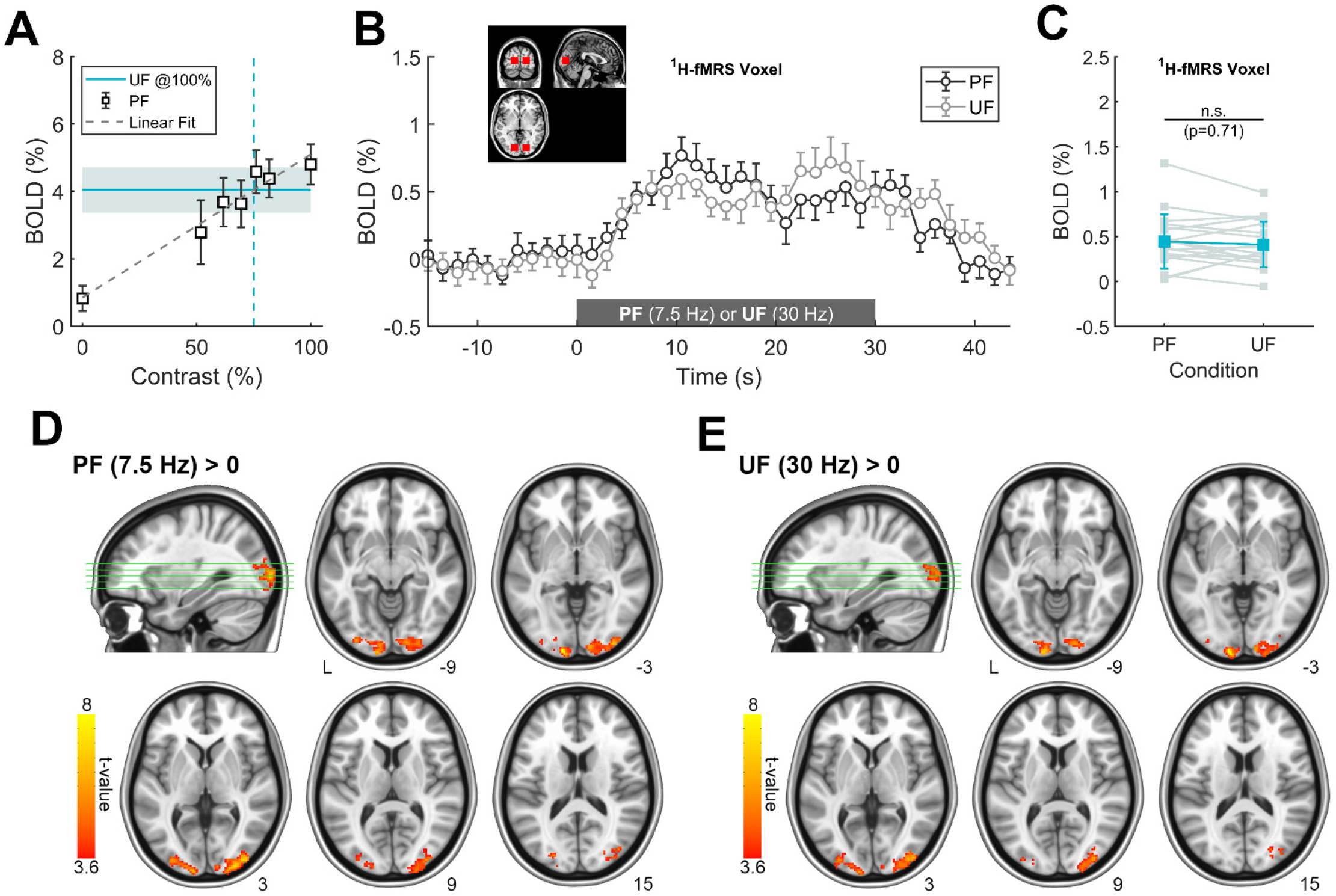
Main effect of stimulation assessed by fMRI analysis. (A) Calibration of image contrast to match BOLD response in V1 to PF and UF obtained in a preliminary session. The contrast of the PF image was reduced to 75% for subsequent stimulations (i.e. common to all subjects). (B) Mean time-course of BOLD signals in the transition between rest and PF or rest and UF, averaged over the fMRI voxels corresponding to the subject-specific spectroscopic VOI. (C) BOLD percent change during the experimental conditions, averaged over the fMRI voxels corresponding to the subject-specific spectroscopic VOI. No statistically significant difference in BOLD response was found between the two conditions (unpaired two-sample t-test, p>0.71). (D,E) Statistical maps for group-averaged positive effect of the visual stimulation (PF and UF) versus rest. Normalized maps are thresholded at p<0.001, with a FDR correction at the cluster level (corresponding to q<0.05), and overlaid on MNI template.

### Different BOLD responses in secondary visual areas to perceived and unperceived flickering

To better characterize the effect of the two different stimulations, we estimated the main effect of the flickering frequency. The main effect of PF appeared in the lateral occipital cortices, but not in V1 (Figure 3B and Table 2). Although the electrophysiological activity in V1 could not be directly assessed in our experiment, based on the literature (27) we can obtain a rough surrogate of V1 output by evaluating fMRI signals in the secondary visual areas, which receive input directly from V1. The average BOLD change in these areas (Brodmann Areas 18 and 19) was significantly higher during PF than UF (0.61±0.29% versus 0.39±0.18%, paired two-sample t-test, p=0.008), while the response in V1 (Brodmann Area 17) was similar for the two stimulations (0.85±0.45% versus 0.80±0.42%, paired two-sample t-test, p=0.72) (Figure 2C,D), indicating a larger output from V1 during PF compared with UF. Thus, V1 exhibited the same BOLD signal despite known differences in visual processing for PF and UF (26).

**Figure 3.**
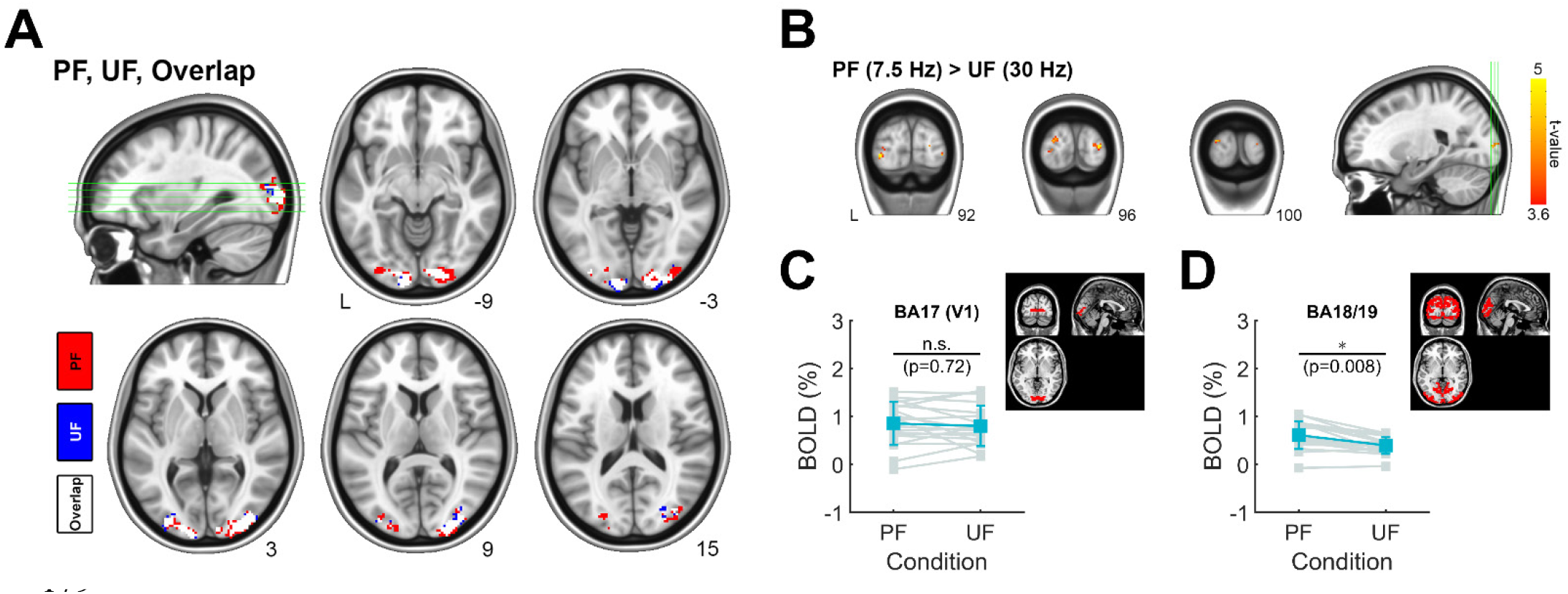
Differential effect of stimulation assessed by fMRI analysis. (A) Overlap between activation maps relative to PF and UF. (B) Differential effect of stimulation (PF>UF). Statistics are thresholded as in Figure 2 (p<0.001, q_FDR_<0.05). The differential response related to perception is localized in lateral occipital cortex (secondary visual areas), with no responding voxels inside V1. For comparison, the inverse differential effect of stimulation (UF>PF) has no significant responding voxels (not shown). (D) BOLD percent change averaged over the fMRI voxels corresponding to the Brodman Area 17 (i.e., V1). Within V1, there is no difference between PF and UF condition (unpaired two-sample t-test, p=0.72). (E) BOLD percent change averaged over the fMRI voxels corresponding to the Brodman Areas 18 and 19 (e.g., including V2, V3a, V4v, V5/MT). Within these areas, the response to PF is significantly larger than the corresponding response to UF (unpaired two-sample t-test, p=0.008).

**Table 2.**
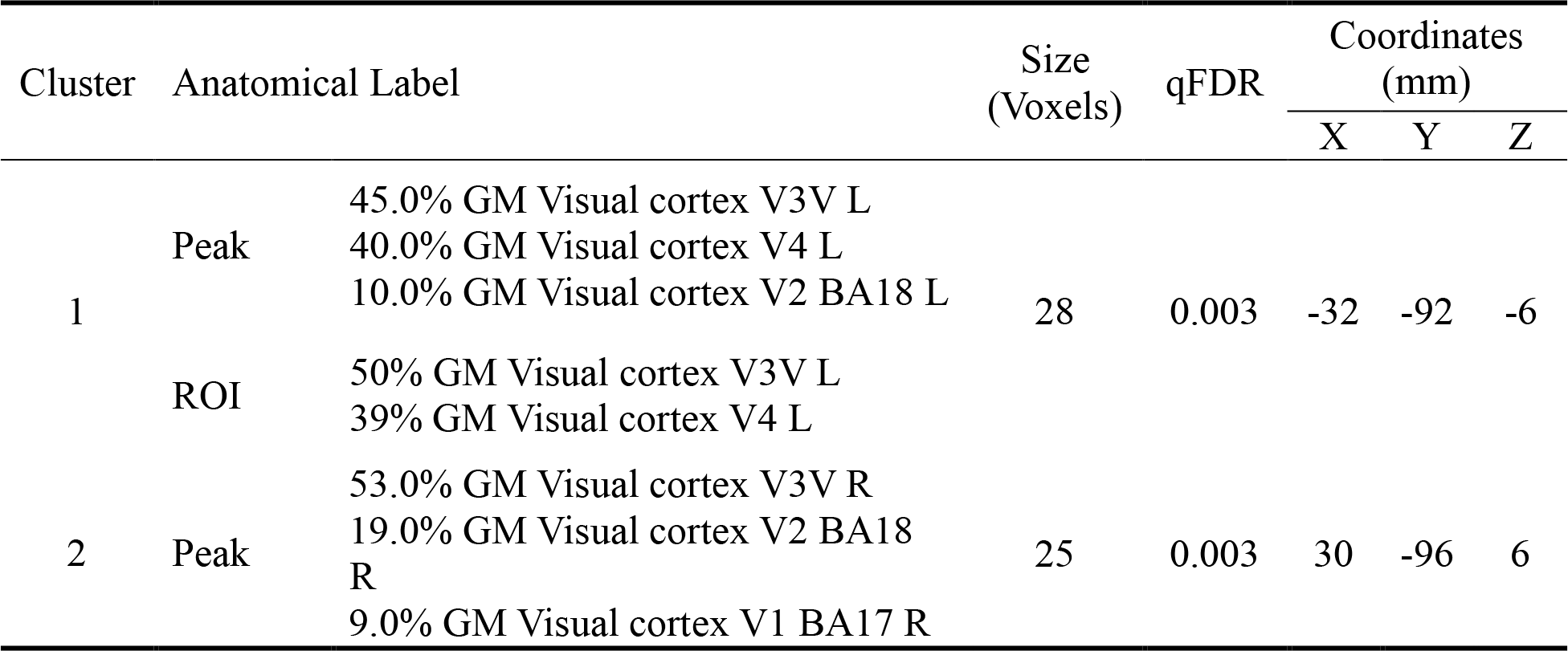
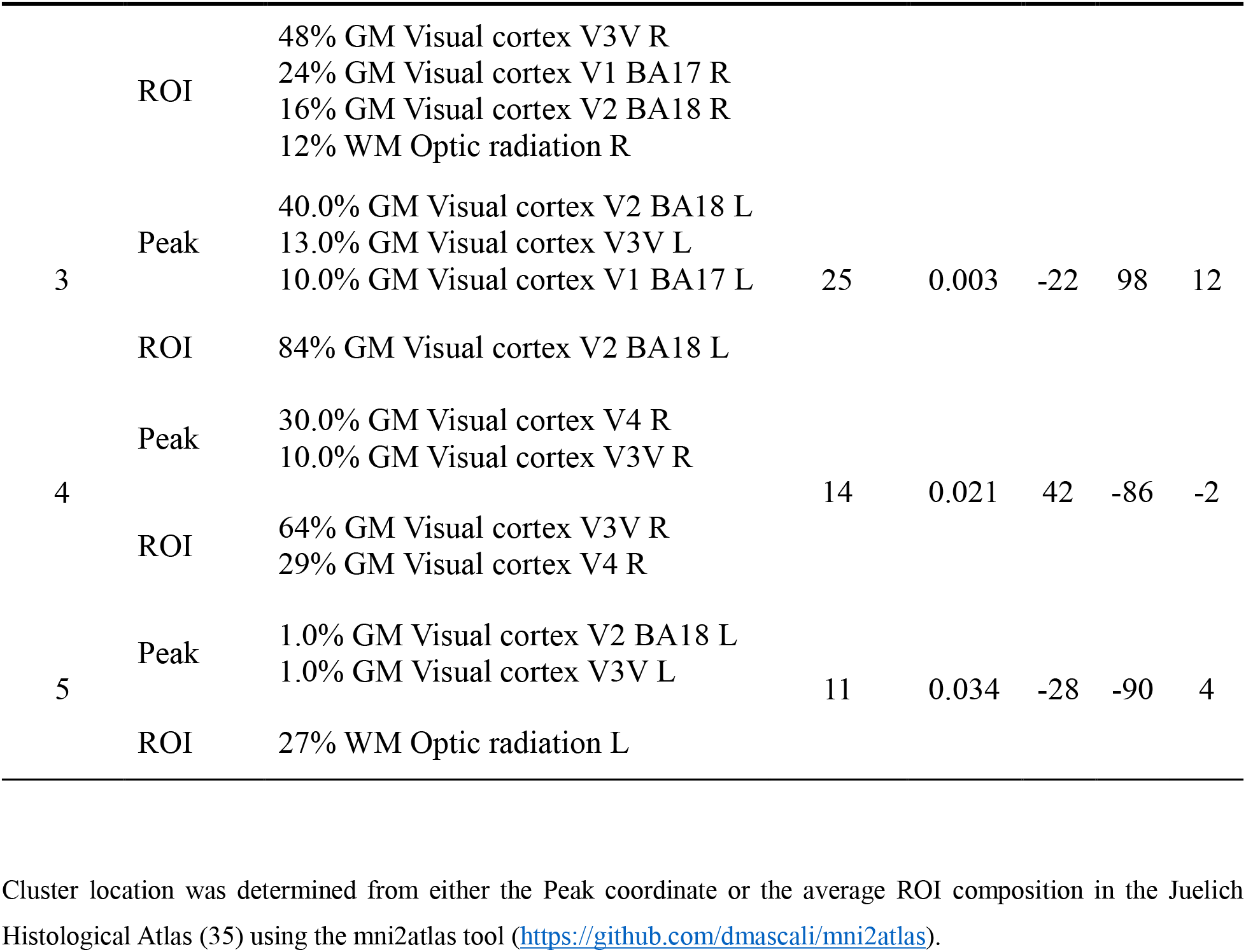
Cortical regions preferentially activated by PF compared to UF.

### Dissociation between metabolic and BOLD responses in V1 during loss of visual perception

To assess whether metabolism was sensitive to stimulus processing within V1 we performed single-voxel 1H-fMRS. The reproducible BOLD response in the occipital lobe elicited by the PF and UF stimulations allowed a very accurate VOI positioning for 1H-fMRS. High quality and artifact-free spectra (water linewidth 7.2±0.6 Hz, Figure 4A) were reliably obtained from almost all subjects (Table 1 and Figure 4 – figure supplement 1). Compared with resting conditions, the lactate and glutamate concentrations within V1 increased by 0.29±0.18 μmol/g and 0.31±0.20 μmol/g, respectively, during the PF stimulation corresponding to an increase of about 28% and 3%, respectively, over the baseline (paired two-sample t-test, q_FDR_=0.001), whereas they both remained at their basal levels (−0.04±0.13 μmol/g, q_FDR_=0.42 for lactate, and 0.03±0.20 μmol/g, q_FDR_=0.63 for glutamate) during the UF stimulation. The lactate and glutamate responses were significantly different (paired two-sample t-test, q_FDR_=0.01 for lactate and q_FDR_=0.003 for glutamate) among the two stimulation conditions (Figure 4B). No other metabolites among those quantified showed a reliable stimulation-dependent change (Table 3). We were unable to detect a reliable change for aspartate (paired two-sample t-test, q_FDR_=0.98).

**Figure 4.**
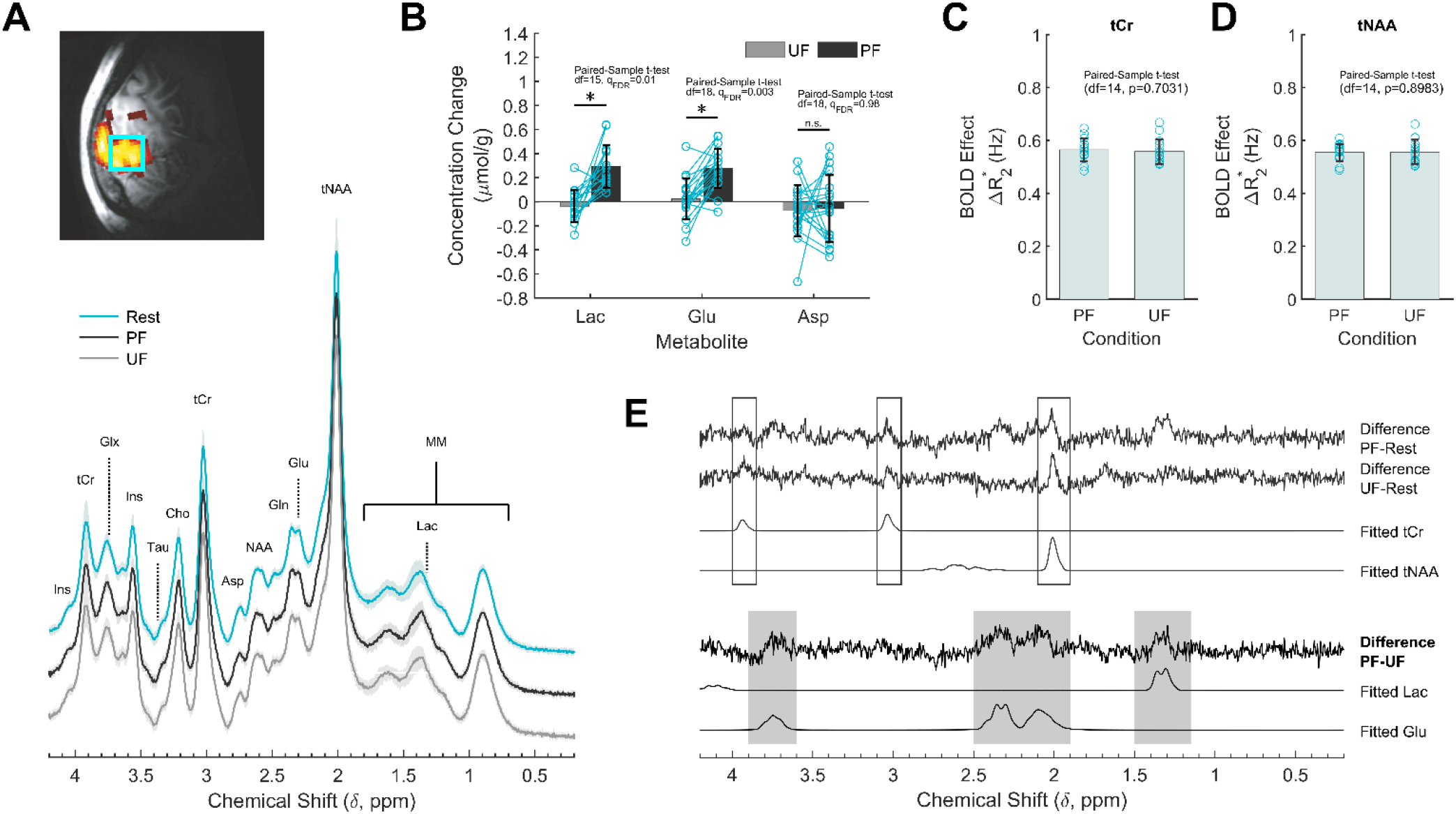
^1^H-fMRS analysis. (A) Spectroscopic data acquired during resting condition (R, cyan) as well as PF (black) and UF (gray) averaged across subjects. A single-subject representative voxel location is reproduced on a parasagittal view of the BOLD activation and superimposed on the anatomical scan from the same subject. For visualization purposes, the processing of the spectra included frequency and phase correction of single transients, averaging, eddy currents correction, and Fourier transform. (B) Lactate, glutamate, and aspartate concentration changes during the stimulation conditions, relative to the rest conditions acquired immediately before. Data are averaged across subjects. There is significant increase in lactate (+28%) and glutamate (+3%) levels induced by PF stimulus, but not by UF stimulus. The concentration changes of the two metabolites were significantly different across the stimulation conditions (q_FDR_=0.01 for lactate and q_FDR_=0.003 for glutamate), while there was no change for aspartate (q_FDR_=0.98). (C,D) Spectral tCr and tNAA linewidth changes induced by the PF and UF stimuli shows no statistically significant difference (p>0.7). (E) Differences between spectra acquired in the three experimental conditions. For reference, the corresponding LCModel fits are reported on the bottom for the Lac and Glu signals. tCr and tNAA singlets showed the expected BOLD related features: there is a difference between stimulation and rest, but the difference spectra between the active conditions are within the noise. In the regions of lactate and glutamate the difference spectra between PF and rest and between PF and UF are similar, while they are clearly disinct from the difference spectra between UF and rest.

**Table 3.** Modulations of metabolic profile of V1 during PF and UF stimulations.

| Metabolite | N | CRLB (%) | ΔConcentration (STIM-REST) | |
| --- | --- | --- | --- | --- |
|  |  | All epochs | UF | PF |
| | | CRLB (%) | mean±SD (μmol/g) | mean±SD (μmol/g) |
| Asp | 19 | 11 | -0.06±0.17 | -0.07±0.26 |
| tCr | 19 | 4 | -0.02±0.06 | 0.00±0.08 |
| GABA | 15 | 16 | -0.12±0.28 | -0.02±0.22 |
| Gln | 19 | 12 | -0.03±0.07 | -0.04±0.10 |
| Glu | 19 | 5 | 0.03±0.20 | 0.31±0.20 (*) |
| GSH | 18 | 14 | -0.03±0.15 | 0.06±0.12 |
| Lac | 16 | 26 | -0.04±0.13 | 0.29±0.18 (*) |
| Ins | 19 | 5 | -0.02±0.09 | -0.01±0.11 |
| NAA+NAAG | 19 | 3 | -0.02±0.12 | 0.00±0.09 |
| tCho | 19 | 7 | 0.02±0.12 | 0.02±0.14 |
| Tau | 16 | 24 | 0.06±0.18 | -0.02±0.17 |

Only metabolites with CRLB<30% in at least 15 subjects are reported (35% for lactate). CRLB are averaged between all the epochs.
(\*): Statistically significant.

To confirm our fMRI result of similar BOLD effect during PF and UF, we examined linewidth narrowing of total creatine (tCr) and total N-acetylaspartate (tNAA) signals. We found a stimulation-induced decrease of tCr and tNAA linewidth during both PF and UF (Figure 4C,D) that was not statistically diffferent between the two conditions (0.56±0.04 Hz vs 0.56±0.05 Hz for tCr, paired-sample t-test, p=0.70; 0.55±0.03 Hz vs 0.56±0.05 Hz for tNAA, paired-sample t-test, p=0.89).

To substantiate our 1H-fMRS result, we determined the difference spectra between conditions, which mainly consisted of uncorrelated noise and only a few correlated residuals (Figure 4E). Specifically, the difference between PF and rest spectra showed a signal in the region corresponding to lactate, and in spectral regions corresponding to glutamate; both signals were absent in the difference between UF and rest spectra. The difference spectra also featured some narrow peaks corresponding to the main singlets of the spectrum, particularly tCr and tNAA, as a result of BOLD-induced line narrowing (36). Similar residuals on tCr and tNAA were recognizable in the difference spectrum between UF and rest, but not in the difference spectrum between the two active conditions, again consistent with the evidence of a similar BOLD effect on spectral linewidth elicited by either of the stimulations. Overall, the only correlated signals that survived in the difference spectrum between PF and UF were lactate and glutamate, which strongly supports the significance of the concentration changes based on LCModel quantifications.

## Discussion

The cortical gray matter of the brain features one of the highest metabolic rates of all organ tissues of the human body. Although energy is recognized as a limiting factor for the human cerebral cortex (37, 38), the increase in lactate concentration occurring upon sensory stimulation isn’t the result of limited oxygen availability (39, 40), as it is for skeletal muscle. Why the cerebral cortex upregulates glycolytic metabolism for sensory information processing is unknown, but it is well-established that glycolysis serves specific neurophysiological and neurobiological purposes, such as axonal vesicle transport, vesicle recycling, action potential waveform modulation, reuptake of neuroactive compounds, and dendritic spine remodeling (reviewed in (31)). Furthermore, lactate is known to be implicated in cognitive processes occurring during waking activity, like learning and memory (41–44), although the exact underlying mechanisms are still debated (45, 46).

In the present study, we asked whether alterations in visual perception are also reflected in metabolic changes within the primary visual cortex in humans. We report that stimulus perception affects the lactate and glutamate response in V1. The PF and UF stimulations elicited, by experimental design, the same average BOLD signals increase inside the spectroscopic voxel, indicating an equivalent degree of neurovascular coupling and possibly of local synaptic activity in the two experimental conditions (47). Yet, we observed a significant increase in the regional lactate and glutamate levels only during the PF stimulus, with no appreciable change during the UF stimulus compared with resting conditions. A temporal dissociation between BOLD and lactate changes has been previously reported during repeated photic stimulations (48). In that case, BOLD response was preserved, but not the lactate and glutamate accumulation, possibly due to habituation of neuronal firing. Metabolic adaptation, in terms of glutamate levels, in the presence of constant BOLD and electrophysiological activity was also reported in epilepsy (49). These results suggest that the physiological mechanisms underlying BOLD signals and energy metabolism do not necessarily overlap under all experimental conditions.

Our results suggest that lactate and glutamate may be dissociated from BOLD changes when cortical input and output are differentially modulated by e.g., intracortical inhibition (26). In fact, an overall switch toward inhibition is expected to reduce the energy request of the brain, thus impacting on metabolic rates independently of BOLD response (2). Accordingly, changes in excitation/inhibition equilibrium have been proposed as a source of variability of the CBF/CMRO_2_ coupling ratio (50). In more general terms, the metabolic response seems capable of differentiating neural states that are intrinsically distinct, although they induce the same BOLD response. Similar BOLD signal changes in correspondence to substantially different alterations in tissue lactate and glutamate level during visual flickering could not be predicted a priori unless assuming fundamental differences in information processing during stimulation. In particular, the firing rate of layer IV neurons that receive input from LGN is higher during UF than PF, as evidenced by the synchronization of these neurons to the stimulus frequency (16–21), yet we found that lactate and glutamate increases during PF, not UF. These findings indicate that the stimulation-induced effect on metabolite concentrations is happening downstream the input stage in layer IV, and specifically during the intracortical processing involving output layers II/III. This conclusion is supported by the fact that high-frequency flickering (30-60 Hz) abolished MUA, a measure of neuronal spiking (putatively from layer II/III) (26), but not single neuron recordings from layer IV (16).

Using compartmentalized metabolic modeling, we have recently obtained evidence that the above-mentioned outcome can be explained by distinct signaling mechanisms underlying spiking and synaptic activity (e.g., pre- and postsynaptic ionic currents) that are indeed frequency-dependent (30, 51). We previously reported that chromatic and achromatic flickering at the same temporal frequency elicited the same neurochemical response in V1 despite differences in the responding neuronal populations (i.e., blob versus interblob) (10). Together with the results that we report here, these observations suggest that lactate and glutamate, and hence aerobic glycolysis, are both sensitive to cortical processing (e.g., input/output or excitation/inhibition balance) rather than the targeted neuronal population. Our results can be interpreted in keeping with the concept that increased glutamate levels reflect an upregulation of the malate-aspartate shuttle (MAS) (52), although we were unable to report significant differences in aspartate levels likely due to insufficient sensitivity of MRS at 3T. Notably, MAS does not necessarily correlate with mitochondrial respiration and cerebral blood flow, as evidenced by the findings that (i) oxidative metabolism and CMRO_2_ are enhanced at both low and high flickering frequencies (26), and (ii) glutamate and aspartate, but not lactate, correlate with BOLD signals during low-frequency (∼8 Hz) visual stimulation (10, 53). To what extent aerobic glycolysis and MAS can dissociate from oxidative phosphorylation in the brain remains to be established, but previously published data strongly indicates that the increase in lactate and glutamate levels is the consequence of the surge in glycolysis and the ensuing accumulation of NADH taking place during neuronal activation (54).

While the similar BOLD response in V1 is a direct consequence of our experimental design, we additionally found that secondary visual cortices were recruited to a larger extent during PF stimulus than UF stimulus, supporting the argument that their activation is necessary for perception (55–57). The preferential response of secondary visual areas to perceived stimuli that we observe here broadly confirms previous results of a strong BOLD activity in ventral (“visual-for-perception” processing) and dorsal (“visual-for-action” processing) streams during visible stimuli and a reduction of their activity in conditions of invisible stimulation (58).

The role of primary visual cortex in stimulus perception is an intense area of research (59–63). Previous fMRI studies investigating the dependence of V1 BOLD activity on visual perception produced controversial results, which might be related to the difficulty of disentangling perception from spatial attention. Indeed, it has been shown that attention, but not perception, modulates the BOLD signal in human V1 (32). In fact, perception was found to enhance the BOLD response within V1 for specific experimental designs (e.g., flash suppression of perception) (64). Our study employed a substantially different stimulation paradigm that specifically allowed us, by adjusting the stimulation contrast, to induce the same BOLD response, and inductively an equivalent mean degree of synaptic activity, within V1 under the two experimental conditions (47). Most importantly, we did not use any additional stimulus (e.g., visual masking) to suppress or modulate visual perception. On the contrary, we used a simple way to modulate perception for extended periods of time (required by 1H-fMRS) while maintaining attention at a nearly constant level, as evidenced by pupillometry and task performance. In particular, we employed the well-known dependence of BOLD response in V1 to flickering, which at full-contrast peaks at 4-8 Hz and settles around 70% of its maximal value even at frequencies above 30 Hz (65), i.e., in the absence of flickering perception. Previous literature reported that the peak activations in V1 and secondary visual areas are stimulation-dependent and occur at distinct temporal frequencies (4 Hz or 8 Hz in V1, and 20 Hz or 40 Hz in secondary visual areas for chromatic or luminance flickering, respectively) (66–70). Interestingly, the preferred frequency of secondary visual areas is just below the CFF for chromatic flickering (∼25 Hz) and luminance flickering (∼50 Hz) (71). In our experiments, the isoluminant chromatic flickering stimulation at 30 Hz is above the CFF and accordingly we found that the BOLD response in secondary visual areas drops substantially compared with the concurrent BOLD response in V1. Overall, by using visual stimulations below and above the CFF and adjusting the contrast of the low-frequency stimulus we were able to modulate perception alongside BOLD activity in secondary visual areas with an unchanged BOLD activity in V1.

Our study has some limitations. For instance, the fMRI measurements have been obtained using 30 s epochs, while the fMRS measurements have been obtained using 4 min epochs. Long acquisition times are required for achieving a good sensitivity of the 1H-fMRS measurements and a reliable metabolite quantification, even at magnetic fields of 3T. Nonetheless, results in both humans (48) and rats (72) have showed that prolonged (i.e., minutes) stimulations elicit a persistent BOLD response with minimal attenuation. Although we cannot exclude a certain degree of neuronal adaptation, we confirmed that the BOLD effect during the fMRS acquisition was indeed relatively stable as determined by the linewidth changes of tCr and tNAA signals. Another limitation to consider is represented by the large volume used for 1H-fMRS measurements. It could be possible that the observed changes in lactate levels include distinct neuronal populations showing non uniform responses. Indeed, although the main differential effect of frequency was located far from the calcarine sulcus, it also involved some mid hemispheric areas presumably included in the spectroscopic voxel. As an illustration, heterogeneity in the response to stimulation within V1 might be due to, e.g., eccentricity (67), which we however minimized by using a 3° foveal stimulation. There is also an hemispherical lateralization for the spatial frequencies (73), but this is not likely to apply to our study, as we used the same radial checkerboard (i.e., including many spatial frequencies) for both PF and UF. The finding that the changes in BOLD signal during PF and UF were nearly double in the anatomical (i.e., Brodmann atlas-based) V1 compared with the spectroscopic VOI indicates a substantial partial-volume effect. The associated loss of sensitivity entails that the lactate concentration change that we report here might in fact be under-estimated. Finally, we cannot exclude that feedback input to V1 from secondary visual areas might be taking place during stimulus perception (74). However, under our experimental design we were able to fully control BOLD response in V1 by only acting on stimulus contrast, without altering the subjective perception of the stimuli, which indicates that the BOLD activity in V1 largely reflected stimulus features.

In conclusion, in this study, we report for the first time that visual perception has a measurable metabolic effect on V1. Our findings imply that BOLD fMRI and 1H-fMRS are complementary techniques, capable of highlight different aspects of neural activation and stimulus processing, e.g., under conditions in which synaptic and spiking activity are partially disentangled due to an altered balance between excitation and inhibition.

Although we focused on the visual pathways, it is conceivable that our results can be translated to other sensory modalities. For example, tactile (75) or auditory (76, 77) stimulations during vegetative state can still activate primary somatosensory cortex (S1) or primary auditory cortex (A1), respectively, in the absence of perception and without the involvement of higher-order associative areas.

We suggest that the cortical metabolic profile might be an indicator of sensory perception, in keeping with the dynamics of lactate and glutamate across the sleep-wake cycle (31, 40, 78, 79). In particular, elevated brain lactate and glutamate levels are associated with wakefulness and memory formation, which naturally require the processing of incoming sensory stimuli, like the control exerted by the central visual pathways for either gating or filtering out behaviorally relevant or irrelevant visual information. In particular, aerobic glycolysis and lactate might reflect cortical information processing and, in turn, intracortical communication, in agreement with the relation between regional metabolic rates of glucose utilization and resting-state network dynamics in the cerebral cortex (80–84).

## Materials and methods

### Setup

MR measurements were performed on a 3T head-only scanner (Siemens Allegra, Erlangen, Germany), equipped with high performance gradients (amplitude 40 mT/m, rise time 100 μs). A custom-built quadrature surface coil (loop and butterfly design, Rapid Biomedical, Rimpar, Germany) was used for both RF transmission and detection. The coil design traded usable volume (see Figure 2 – figure supplement 1) for peak sensitivity. For imaging, first and second order shim terms were adjusted using the automatized Siemens routine based on field map acquisition and fitting. For MRS, shimming was optimized using FASTMAP with EPI readout (85) and manually refined when necessary to reach a water linewidth of less than 9 Hz.

### Subjects

A total of 32 healthy volunteers were initially enrolled for this study after they gave informed consent, according to the Helsinki declaration and to European Union regulations, and following the approval by the Ethics Committee of Fondazione Santa Lucia (Rome). Exclusion criteria included any kind of previous neurological or psychiatric disease and impaired visual acuity. Thirteen subjects were discarded either upon online fMRI processing (see below) or in early quality control on data, because of motion (3) or suboptimal anatomical features, with the most activated area in an unfavourable position respect to the coil sensitive volume (10). Nineteen healthy volunteers (10 females, 9 males; age 25±4 years, mean±SD; age range 20 to 35 years) were thus considered for this study. Sample size calculations performed before the study assumed a two-tail paired t-test design, a power of 0.95 and were based on an effect magnitude for lactate change (photic stimulation vs rest) of 0.20 ± 0.15 μmol/g as estimated in our previous works at 7T (10). Reduced sensitivity of 3 vs 7T was epmpirically accounted for by larger VOI and extended data averaging (144 transients per condition, 10 ml VOI at 3T vs 64 transients per condition, 8 ml VOI at 7T). The resulting required sample size of 10 was roughly doubled to account for multiple comparisons and other suboptimal procedures at 3T compared to 7T.

### Visual stimulation

Visual stimulation consisted in a radial checkerboard flickering either at 7.5 Hz (perceived flickering stimulation, PF) or at 30 Hz (unperceived flickering stimulation, UF). The alternating frames included either a gray-green or a gray-red checkerboard. The stimuli were programmed in Cogent 2000 version 1.29 working under Matlab 2006b (The Mathworks, Natick, MA, USA). and delivered using an MR-compatible fMRI system with stereo 3D goggles (VisuaStim Digital, Resonance Technology, Inc., Northridge, California, USA). Subjects were fitted with the VisuaStim video goggles (Resolution: SXGA 1280×1024 pixels, Refresh Rate: 60Hz, Field of View: 30 degrees Horizontal x 24 degrees Vertical, White Luminance: 70cd/m2 max, Contrast Ratio: intrinsic 100:1 measured per VESA FPDM Standard).

### Experimental protocol

Visual stimulations were presented in eight (fMRI) or four (fMRS) epochs, with stimulation epochs (either PF or UF) interleaved by rest (isoluminant, uniform grey images) epochs. Since the CFF for loss of perception is higher for luminance than for chromatic flickering (25), before each session the brightness of green squares during the UF condition (i.e. isoluminant condition) was adjusted interactively by the subject, who piloted increasing and decreasing brightness ramps and was instructed to identify the brighness levels corresponding to loss of luminance flickering perception of the resulting yellow. The green level was then set midway between the two perceptual vanishing levels. The stimulus contrast was adjusted in preliminary acquisitions on five subjects, in order to induce a comparable BOLD response to PF and UF in V1. During the initial fMRI sessions epochs lasted 30 seconds each (total 4 minutes), while for fMRS acquisitions epochs were 4 minutes long (total 16 minutes). Each subject underwent 1 fMRI and 2 fMRS runs (10 minutes apart, without moving the subject from inside the scanner); the order of PF and UF conditions was counterbalanced within each subject, and the initial stimulation type was randomized between subjects.

### Task

In order to maintain visual fixation and keep the attentional state constant, the subjects were asked to focus on a central target (a cross) and to press a button whenever the target rotated. Subjects were specifically instructed to maintain their attention on the fixation cross rather than focusing on reaction times (i.e., speed to push the button). The number of rotations was constant across the different epochs (3 for each fMRI epoch and 24 for each 1H-fMRS epoch, or approx. 1 rotation every 10 seconds), while the exact timing of the rotation was pseudo-randomized (range 2-18 seconds).

### Anatomical and fMRI data acquisition

Each study started with an anatomical acquisition (MPRAGE T1-weighted volumetric scan, resolution 1.2×1.2×1.2 mm^3^, para--axial slices, in-plane FOV 190×70 mm^2^, TE=4.38 ms, TR=2000 ms, TI=910 ms, FA=8°). Then, one fMRI session (pseudo-randomized order of stimulation across subjects) was acquired (gradient echo with EPI readout, resolution 2.2×2.2×2.2 mm^3^, 26 para--axial contiguous slices, FOV 190×70 mm^2^, TE=30 ms, TR=1500 ms, FA=70°). fMRI scans were processed online for subsequent MRS voxel positioning (online processing included motion correction, smoothing, cross-correlation with a square-wave model; the two scans following each condition change were discarded in order to reduce the effects of BOLD signal transients. Online processign was discarded after voxel positioning).

To confirm the absence of any detectable brain pathology in our subjects, T1 and T2 weighted anatomical scans were acquired with a standard volume birdcage coil after the end of the functional scans. Anatomical scans included an MPRAGE acquisition (resolution 1.0×1.0×1.0 mm^3^, para-axial slices, in-plane FOV 256×160 mm^2^, TE=2.48 ms, TR=2150 ms, TI=1000 ms, FA=8°), that was later used during the post-processing for normalization purposes.

### fMRS data acquisition

The spectroscopic voxel (size 25×20×20 mm^3^) was localized in the most activated area within V1, based on both anatomical scan and results of the online fMRI processing. The voxel was located either left or right of the interemispheric fissure to minimize the cerebrospnial fluid fraction in the VOI. Two MRS sessions were acquired with an optimized, in-house written STEAM sequence (TE=7 ms, TM=50 ms, TR=3000 ms, FA=70°) which included outer volume saturation and VAPOR water suppression (86, 87). An eight-step phase cycle was used; transients were averaged within each phase cycle, and each phase cycle was saved separately for further processing. Water unsuppressed data were acquired from the same voxel for eddy currents compensation (88). In order to minimize T1 weighting, the flip angle was kept below the calculated Ernst angle in both fMRI and fMRS acquisitions.

### Pupillometry

In order to monitor attentional state with a physiological parameter, we acquired pupillometry data using an eye-tracking system (Applied Science Laboratories, model 504) equipped with remote pan/tilt optic infrared module and a video camera that was custom-adapted for use in the scanner. Subject gaze position and pupil size data were processed as previously described (33).

### fMRI data processing

fMRI (offline) processing was performed with routines from SPM12 (Wellcome Trust Centre for Neuroimaging, UCL) working under Matlab 2018b, AFNI (89) and FSL5 (90), and with custom Matlab routines. For volume-of-interest (VOI) based analysis, fMRI data were realigned to their mean image. For voxel-based analysis, data were also normalized to the MNI template by using the non linear transform calculated on the MPRAGE acquired with the volume coil, after a linear co-registration that used the surface coil MPRAGE image as intermediate step to best match the volume coil MPRAGE to the fMRI series. A Gaussian smoothing with kernel 4×4×4 mm^3^ was then applied before fitting to a model that included the hemodynamic response. Only for visualization purposes, MR images were assembled with standard image processing tools, without any kind of adjustment. Head motion during fMRI acquisitions was evaluated using the framewise displacement, which was calculated as the L1-norm of the realignment-derived parameters after converting angles to linear displacements (91).

### fMRS data processing

MRS data were preprocessed using jMRUI 5.2 (92) and custom Matlab routines. Data were corrected for residual eddy currents, individually phased and frequency shifted to compensate B_0_ drifts, and averaged in blocks corresponding to each rest or stimulation epoch. The first 8 transients of each epoch, i.e. the first full phase cycle (24 s) were discarded to avoid metabolic transients (3). Subsequent phase cyles were inspected individually. They consistently showed good water suppression and no trace of lipidic contamination. A few 8-transient spectra (maximum one in each epoch) featured anomalous line broadening, line splitting or otherwise reduced quality, putatively related to subject motion or deep inspiration, and were discarded before averaging. Each epoch spectrum was thus the average of 64-72 transients. The resulting averages were finally quantified using LCModel 6.3-1 (93) with a tailored basis set. Basis metabolites included alanine, aspartate (Asp), creatine (Cr), γ-Aminobutyric acid (GABA), glucose, glutamine (Gln), glutamate (Glu), glycine, glycerylphosphorylcholine, glutathione (GSH), lactate (Lac), *myo*-inositol (Ins), *N*-acetylaspartate (NAA), *N*-acetylaspartylglutamate (NAAG), phosphocholine, phosphocreatine, phosphorylethanolamine, *scyllo*-inositol, and taurine (Tau). Glucose, an important marker of energy metabolism, whose changes have also been reported in previous 7T studies (3, 9), was not included in the basis set due to highly unreliable quantification observed in preliminary tests. Metabolite spectra were simulated using GAVA (94), including information on the sequence pulse program. The basis set included also a subject-specific macromolecular (MM) signal, that was acquired on each subject in the occipital region, using a double inversion recovery approach (STEAM, TI1=1700 ms, TI2=520 ms, TE=7 ms, TM=50 ms, TR=2000 ms, FA=90°) (95), that resulted in almost complete metabolite nulling, averaged between subjects, and then modeled with Hankel-Lanczos singular value decomposition. LCmodel quantifications with Cramér-Rao lower bounds (CRLB) above 30% were discarded, except for Lac for which the threshold was set at 35%. Since this study is focused on epoch to epoch metabolic changes, absolute quantification with water referencing was not performed to avoid the associated uncertainty. Metabolites were rather referred to the internal creatine signal, assumed to be 7.5 μmol/g in the VOI. Finally, concentrations measured in homologous epochs were averaged, obtaining for each subject four concentrations, corresponding to two stimulation conditions and to the relevant rest reference (the resting epoch immediately successive to a condition). Eleven metabolites were quantified in at least 15 subjects (80% of participants). These included aspartate, total creatine, GABA, glutamate, glutamine, glutathione, lactate, *myo*-inositol, N-acetylaspartate, total choline, taurine. In order to exclude BOLD adaptation during the 4-min duration of the fMRS epochs, we determined the BOLD effect during fMRS scanning as the kernel size (in Hz) that minimized the amplitude of the difference spectra between stimulated epochs (either PF and UF) and the preceding resting epoch. All spectra were then averaged according to three categories: rest, PF and UF conditions, and spectral differences were calculated between conditions.

### Statistics

For pupillometry and task performance results, statistical comparisons were made using Student’s t-test and One-Way ANOVA on the rest, UF, and PF conditions. No post-hoc test was necessary.

For fMRI results, correction for multiple comparisons in functional voxel-based analysis was performed using False Discovery Rate (FDR) correction. Resulting clusters were also checked through Monte Carlo Simulation using the AFNI tool Alphasim (89) after estimation of residuals smoothness.

For fMRS results, statistical analysis was restricted to those reliably quantified metabolites associated with energy metabolism that showed consistent funcional changes in previous fMRS studies (4, 9, 10), namely Lac, Glu, and Asp. Metabolite concentration changes referred to the corresponding resting epoch and between different active conditions were tested using paired two-sample t-tests, with FDR correction for 9 multiple comparisons.

Data were presented as the mean ± standard deviation (SD). A p-value, or a q_FDR_-value where relevant, of less than 0.05 was considered as statistically significant.

## Acknowledgments

The authors wish to thank Edward J. Auerbach for the FASTMAP implementation on Siemens platform, provided by the University of Minnesota under a C2P agreement, and for his help with the setup of offline shim currents calculation. Siemens Healthineers is acknowledged for providing source code and information on the shim coils.

## Data and materials availability

The study was developed using SPM12 (https://www.fil.ion.ucl.ac.uk/spm/software/spm12/), LCmodel (http://s-provencher.com/lcmodel.shtml), jMRUI (http://www.jmrui.eu/) and AFNI (https://afni.nimh.nih.gov/).

Data used for all the figures and for Tables 2-3 is available as source data to each element. Source data include also custom Matlab code for processing related to each figure.

The raw data include sensitive data. The raw dataset cannot be made available in a public repository because of constraints originally set by the Ethics Committee and included in the informed consent signed by participants. Raw data that support the findings of this study are available from the corresponding author upon signing a MTA that would include:

- A list of authorized researchers.
- A commitment to not disclose the raw data to persons not included in the list.
- A commitment to destroy the raw data when legitimate use is finished. Commercial use of the raw data is not allowed.

## Ethics statement

All experiments with human subjects performed by the authors complied with all applicable ethical standards, including the Helsinki declaration and its amendments, institutional/national research committee standards, and international/national/institutional guidelines.

## Supplementary Materials

**NOTE**

Following eLife style suggestions, Supplementary figures are merged to main text as “Figure supplements” and linked to main figures as indicated by the title of each supplementary figure

**Figure 1 - figure supplement 1.**
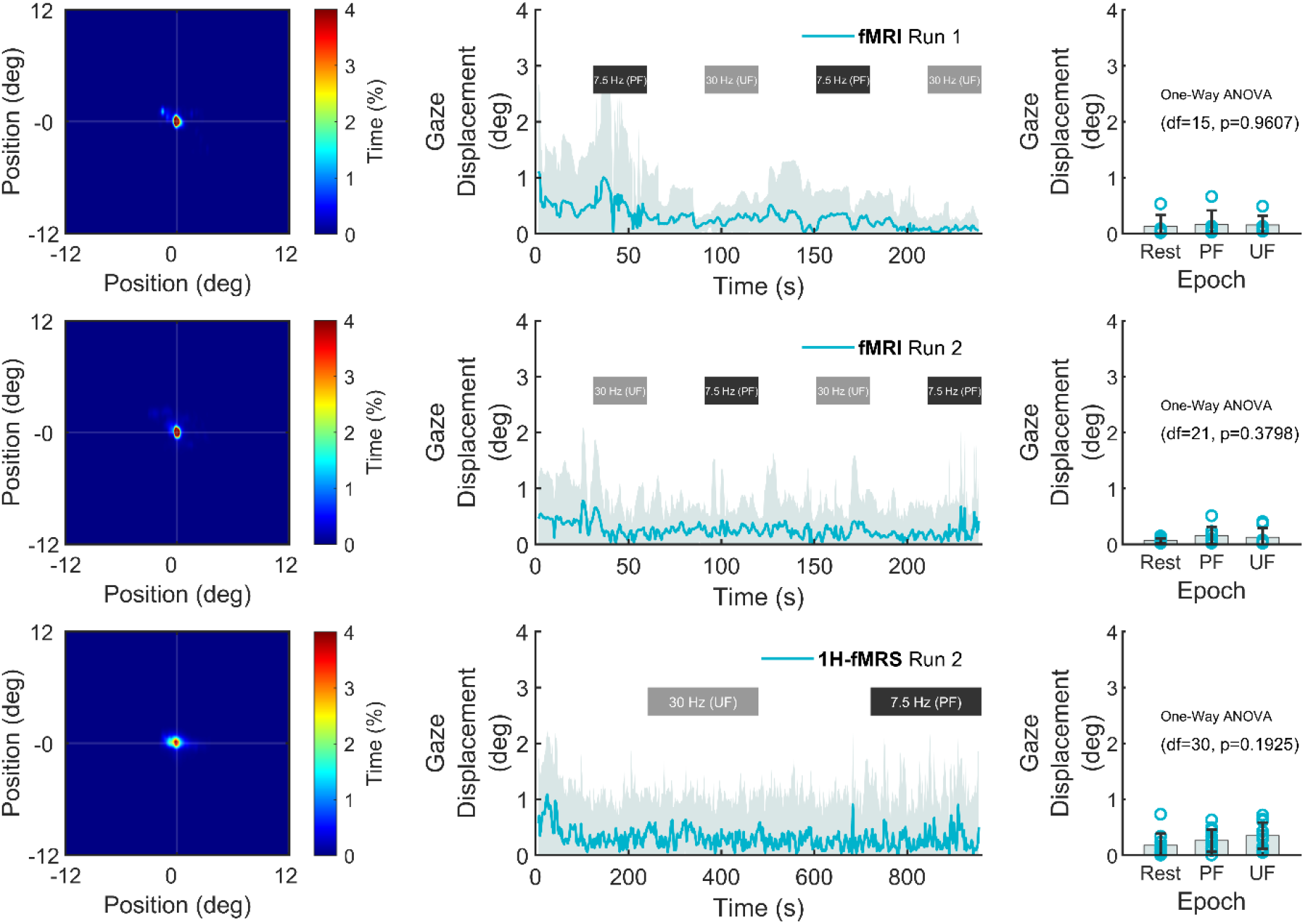
Left: average heatmap of eyes position (across subjects) during the different sessions. (Center) Stability of mean gaze displacement from the fixation point. Right: gaze displacement was not different across conditions (One-Way ANOVA, p>0.19). Error bars correspond to SD. Sessions are shown in different rows (Top: fMRI Run 1; Middle: fMRI Run 2; Bottom: 1H-fMRS Run 2).

**Figure 1 - figure supplement 2.**
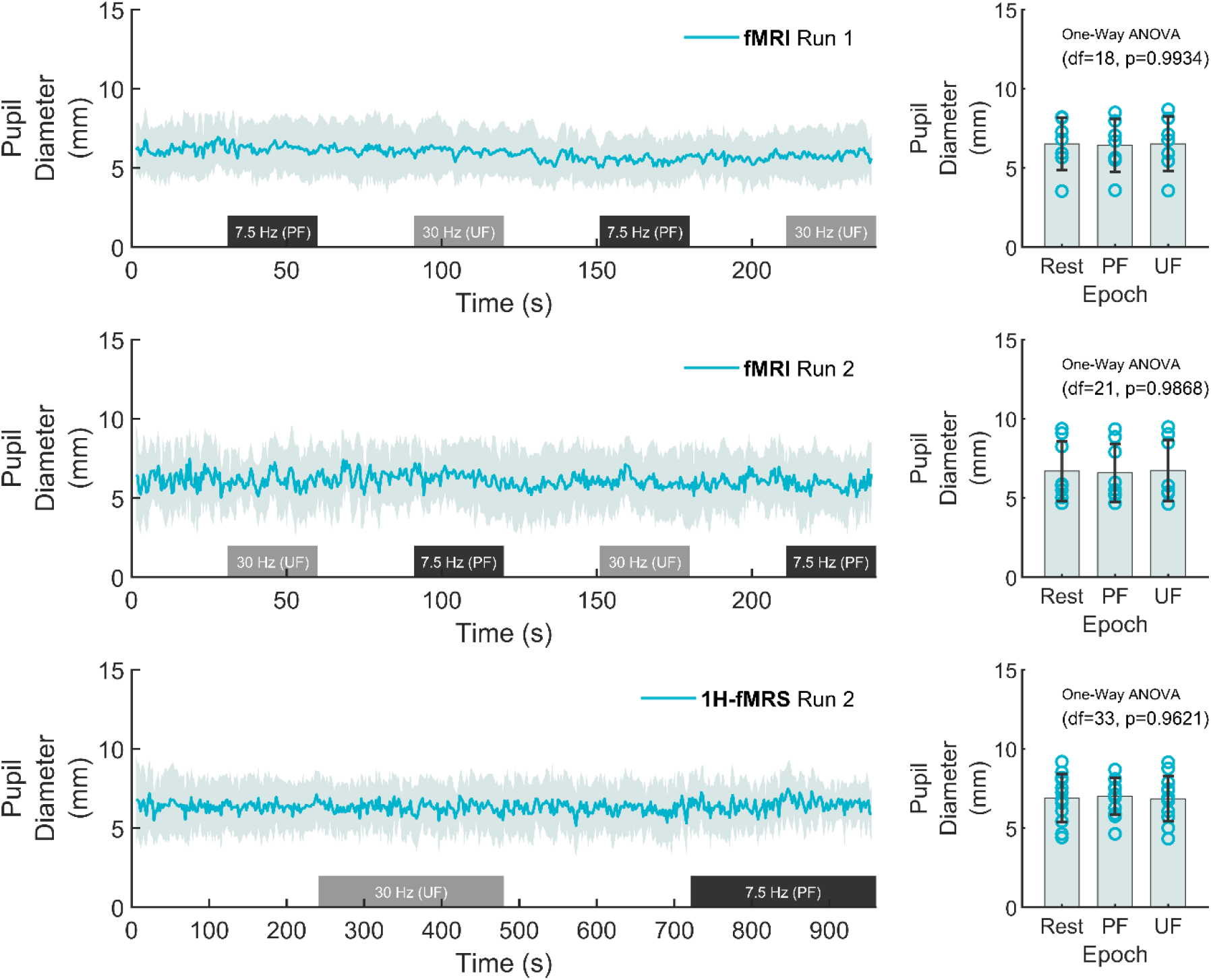
Left: average pupil diameter (across-subjects) during a representative session (1H-fMRS Run 1). Right: mean pupil diameter was not statistically different across conditions (One-Way ANOVA, p>0.96). Error bars correspond to SD. Sessions are shown in different rows (Top: fMRI Run 1; Middle: fMRI Run 2; Bottom: 1H-fMRS Run 2).

**Figure 1 - figure supplement 3.**
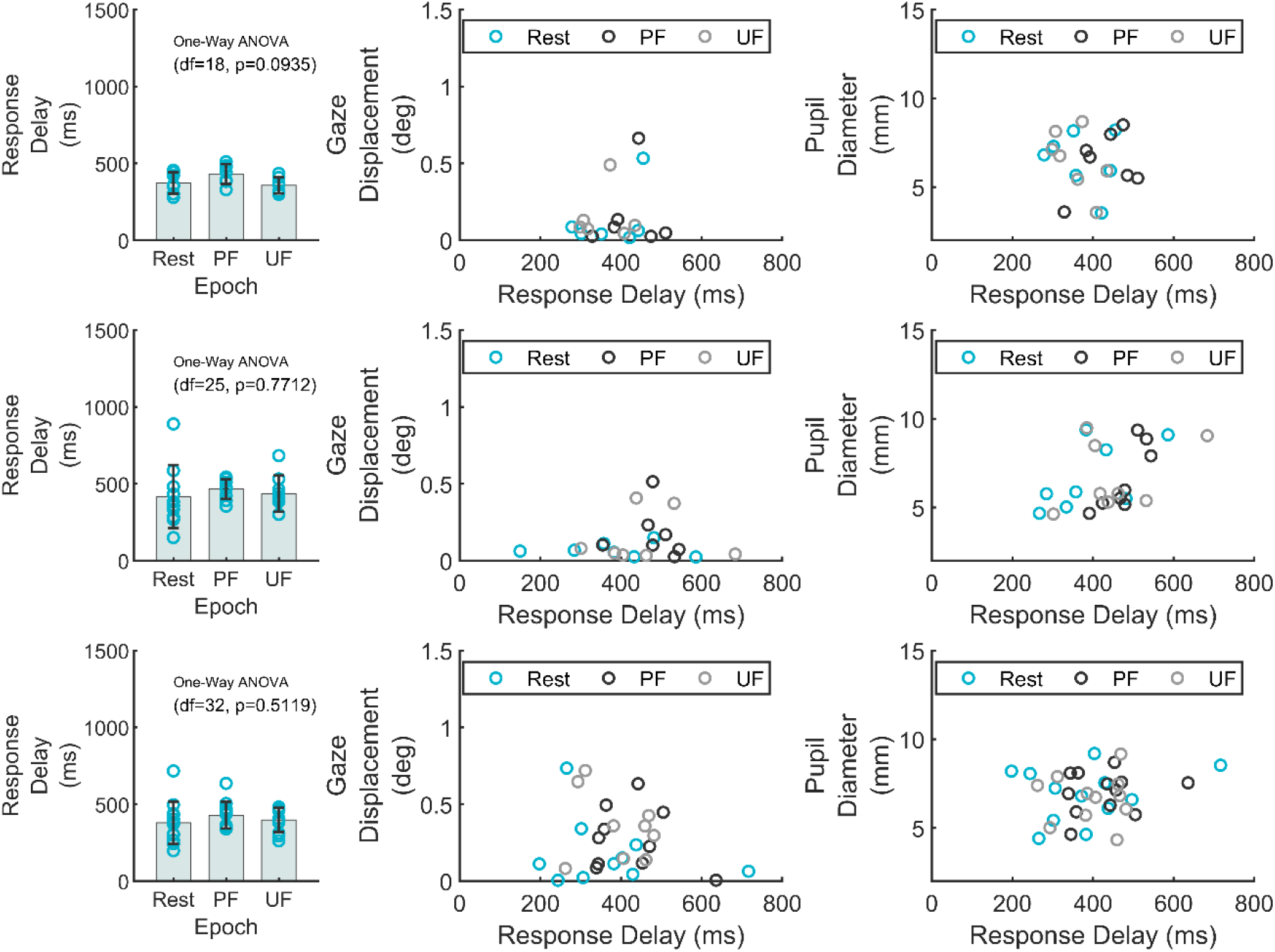
Left: task performance in terms of response delay was not statistically different across conditions (One-Way ANOVA, p>0.09). Error bars correspond to SD. Center: there was no correlation between task performance and mean gaze displacement (r^2^<0.06, p>0.47). Right: there was no correlation between task performance and mean pupil diameter (r^2^<0.08, p>0.49). Sessions are shown in different rows (Top: fMRI Run 1; Middle: fMRI Run 2; Bottom: 1H-fMRS Run 2).

**Figure 2 - figure supplement 1.**
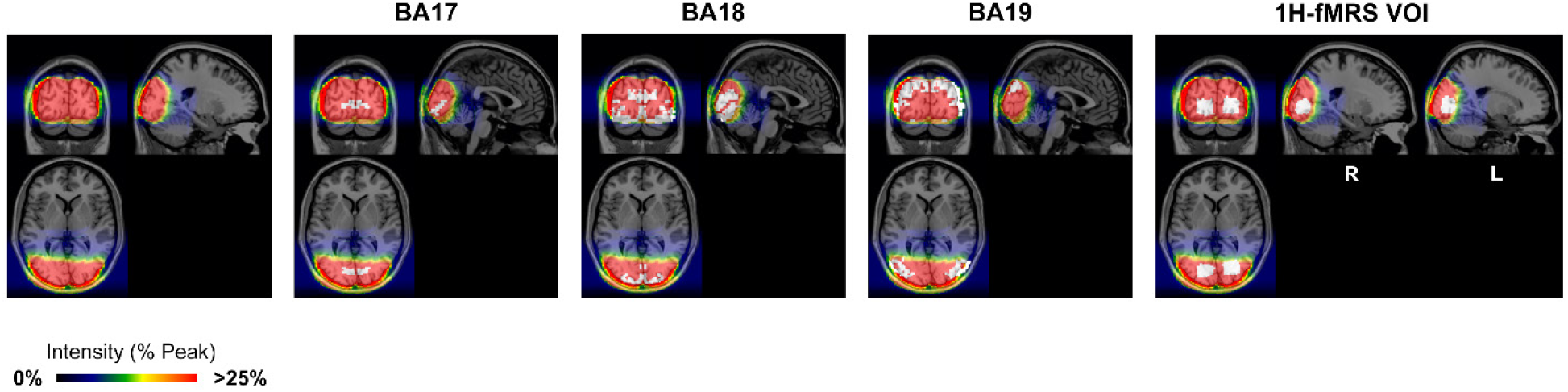
From Left to Right: intensity map of EPI images acquired with the surface coil normalized to MNI template, superimposed to different ROIs (in white): BA17 (i.e., V1), BA18, BA19, and the average spectroscopic voxel (1H-fMRS VOI).

**Figure 4 - figure supplement 1.**
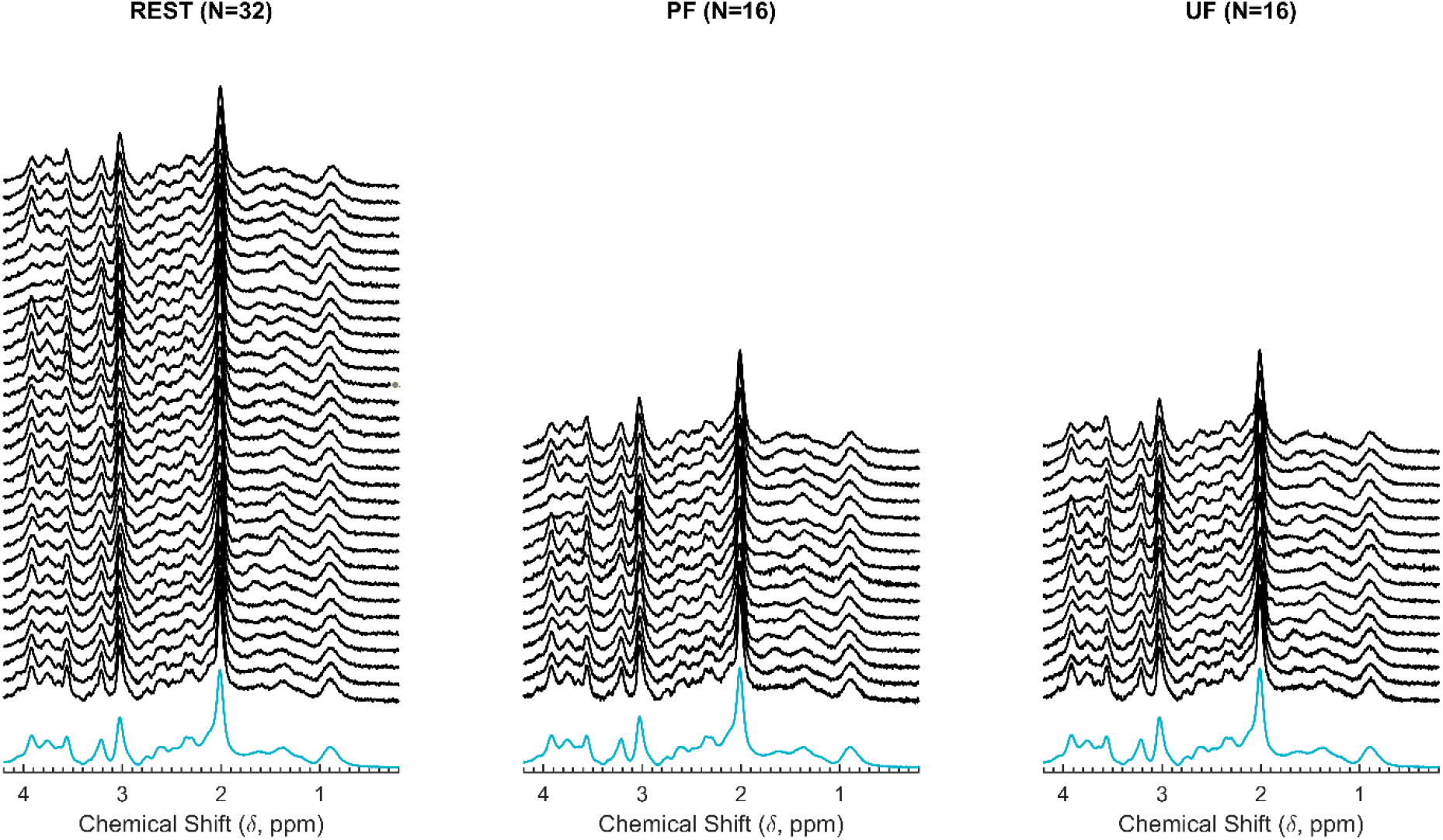
Averaged 1H-fMRS spectra across individual epochs (Left: REST; Center: PF; Right: UF) of the subset of subjects (N=16) who had a reliable quantification for both Lac and Glu. <INSERT Table S1 here followed by a page break>

### Movie S1 (separate file)

The movie shows the pupil of a representative subject during a stimulation cycle including UF and PF epochs, as well as the relevant physiological and behavioral responses.

## Footnotes

1 [Note for reviewing only] Following stile suggesiton of eLife, supplementary figures are linked as children to a main figure. Figure supplements and the relevant captions are at the end of this file.

## Notes

### Competing Interest Statement

The authors have declared no competing interest.

